# Proteomic mapping reveals dysregulated angiogenesis in the cerebral arteries of rats with early-onset hypertension

**DOI:** 10.1101/2022.11.07.515426

**Authors:** Joakim A. Bastrup, Thomas A. Jepps

**Affiliations:** Department of Biomedical Sciences, University of Copenhagen, Denmark

**Keywords:** Hypertension, angiogenesis, cerebrovascular disease, cerebral arteries, DIA-MS

## Abstract

Hypertension is associated with presence of vascular abnormalities, such as remodeling and rarefaction. These processes play an important role in cerebrovascular disease development, however, the mechanistic changes leading to these diseases are not well characterized. Using data-independent acquisition-based mass spectrometry analysis, we determined the protein changes in cerebral arteries in pre- and early-onset hypertension from the spontaneously hypertensive rat (SHR), a model that resembles essential hypertension. Our analysis identified 125 proteins with expression levels that were significantly up- or downregulated in 12-week old SHRs compared to normotensive Wistar Kyoto rats. Using an angiogenesis enrichment analysis, we identified a critical imbalance in angiogenic proteins, promoting an anti-angiogenic profile in cerebral arteries at the early-onset of hypertension. In a comparison to previously published data, we demonstrate that this angiogenic imbalance is not present in mesenteric and renal arteries from age-matched SHRs. Finally, we identified two proteins (Fbln5 and Cdh13), whose expression levels were critically altered in cerebral arteries compared to the other arterial beds. The observation of an angiogenic imbalance in cerebral arteries from the SHR reveals critical protein changes in the cerebrovasculature at the early-onset of hypertension and provides novel insight into the early pathology of cerebrovascular disease.

## Introduction

Many factors are involved in the development of hypertension, which is primarily associated with an increase in the total peripheral resistance, advocating for the presence of vascular abnormalities. Such abnormalities include structural changes to the arterial wall (remodeling and rarefication)^1^, altered excitation-contraction coupling, and changes in the neurogenic and humoral signaling^2^. In hypertension, a general narrowing of the large resistance arteries through to the precapillary microcirculation is associated with eutrophic and hypertrophic remodeling of the arterial wall in most vascular beds^3–5^. We, and others, have demonstrated that arterial remodeling occurs in certain systemic arteries soon after the onset of hypertension (12-weeks old) in the spontaneously hypertensive rat (SHR)^6^, a model that is considered to resemble the main features of essential hypertension in humans without confounding lifestyle and environmental factors^7^. Although the remodeling processes are a key feature of hypertension, the mechanistic pathways involved are not understood fully.

In-depth quantitative proteomics can achieve unique insight into complex biological mechanisms and pathologies^8,9^. We have previously optimized a label-free proteomic workflow for analyzing resistance arteries in the SHR, resulting in the identification of more than 4700 unique proteins^7^. Using gene-overrepresentation analysis, these proteins provided novel mechanistic insight into different biological pathways that were altered during the initiation of systemic arterial remodeling, such as changes in the extracellular matrix (ECM)^7^.

Cerebrovascular diseases, such as stroke, cognitive decline, and vascular dementia are highly associated with hypertension^10–12^. There is evidence that the media-to-lumen ratios of cerebral arteries from the SHR do not display the same level of remodeling as systemic vessels^13^; however, the link between hypertension and cerebrovascular diseases suggests there is a profound effect of hypertension on cerebral arteries^14^. Thus, the aim of this study was to uncover protein changes and mechanistic pathways that are altered in cerebral arteries from hypertensive rats. By determining the hypertension-induced dysregulated pathways in cerebral arteries, we can better understand how hypertension increases the risk of cerebrovascular diseases. Using label-free in-depth proteomic profiling, we reveal that the cerebral arteries from SHRs with early-onset hypertension have several dysregulated proteins associated with angiogenesis, which is not seen in mesenteric or renal arteries of age-matched SHRs.

## Methods

### Experimental animals

The animal experiments were approved by local Animal Care and Use Committees. Experiments were performed in accordance with the directives of the Danish National Committee on Animal Research Ethics, and Danish legislation on experimental animals. Rats were made unconscious by a single percussive blow to the head, in accordance with the methods of killing animals described in annex IV of the EU Directive 2010/63EU. Directly after the onset of unconsciousness, cervical dislocation was used to euthanize the rats. Cohorts of male SHRs (SHR/KyoRj) and WKYs (WKY/KyoRj) at 6 and 12 weeks of age were group housed and supplied with *ad libitum* water and food access. Rats were kept on a 12 h/12 h light/dark cycle and were transferred to clean cages regularly.

### Dissection of cerebral arteries

Following cervical dislocation, the brain was excised delicately and placed in cold physiological salt solution (121 mM NaCl; 2.8 mM KCl; 1.2 mM KH_2_PO_4_; 1.2 mM MgSO_4_; 25mM NaHCO_3_; 1.6 mM CaCl_2_; 0.03 mM EDTA; 5.5 mM D-glucose) saturated with carbogen (O_2_ 95%; CO_2_ 5%) at pH 7.4. The circle of Willis with primary cerebral arteries (posterior, anterior and middle cerebral arteries) were collected in 1.5 ml Lobind centrifugation tubes (Eppendorf), snap frozen in liquid nitrogen and stored at -80 °C.

A small segment (2-3 mm) of basilar artery from a subgroup of rats was embedded in Tissue-Tek OCT (Sakura) for sectioning and staining.

### Staining and microscopy imaging

Arterial cross sections were collected using a cryostat microtome (Leica CM3050 S). Sections were cut at 12 μm thickness and attached to Superfrost Plus glass slides (VWR) and stored at −80 °C. Arterial sections were stained using Sirius red for connective tissue, as reported previously^7^. Slides were scanned using a ZEISS Axioscan 7 slide scanner with 20×/0.8 Plan-Apochromat objective lens (Zeiss). Images were cropped to individual arterial cross sections and analyzed, blinded, in ZEN (v3.2, blue edition) software. Media and lumen diameters were measured using a profile ruler tool to calculate the media-to-lumen ratio.

### Protein isolation and quantification

The cerebral arteries were homogenized in 100 μl of ice-cold lysis buffer (50 mM Tris pH 8.5, 5 mM EDTA pH 8.0, 150 mM NaCl, 10 mM KCl, 1% NP-40 and 1× complete protease inhibitor cocktail (Roche)) by three rounds of chopping the tissue using dissection scissors and a handheld homogenizer^7^. Medial prefrontal cortex regions (0.3 mm × 0.3 mm × 0.3 mm) were homogenized in 150 μl lysis buffer. Homogenates were centrifuged at 11,000g for 10 min at 4 °C to obtain the supernatant. Protein quantification of the tissue extracts was determined by bicinchoninic acid assay (BCA) (Thermo Scientific).

### Sample preparation for mass spectrometry analysis

Digestion buffer containing 0.5% sodium deoxycholate (SDC) in 50 mM triethylammonium bicarbonate (TEAB) was added to the homogenized tissue extracts followed by heat-treatment for 5 min at 95 °C. Samples were cooled on ice and prepared by filter-aided sample preparation^15^ using flat spin filters (Microcon-10kDa). Samples were reduced and alkylated in digestion buffer containing 1:50 (v:v) tris(2-carboxyethyl)phosphine (0.5 M, Sigma) and 1:10 (v:v) 2-chloroacetamide (Sigma) for 30 min at 37 °C. Samples were digested overnight at 37 °C with 1 μg Typsin/LysC mix (Promega) and 0.01% ProteaseMAX (Promega). Centrifugation at 14,000g for 15 min was used in between the different steps. Peptides were desalted using stage-tips containing a Poly-styrene-divinylbenzene copolymer modified with sulfonic acid groups (SDB-RPS; 3M) material, as described previously^7^. In brief, samples were mixed with 5x volume 99% isopropanol (Sigma) / 1% trifluoroacetic acid (TFA) (Sigma), loaded to the stage-tips, and centrifuged at 1,500g at 4 °C until passed through the SDB-RPS filter. Each filter was washed twice in 99% isopropanol (Sigma) / 1% TFA (Sigma) and 0.2% TFA, respectively. Samples were eluted using 80% acetonitrile (Sigma) / 2% Ammonia (Sigma). After vacuum centrifugation, samples were resuspended in 2% acetonitrile (Sigma) / 0.1% TFA (Sigma).

### Data acquisition by DIA-based mass spectrometry

Samples were analysed on a Bruker timsTOF Pro mass spectrometer (Bruker Daltonics) in the positive ion mode with a Captivespray ion source on-line connected to a Dionex Ultimate 3000RSnano chromatography systems (Thermo Fisher). Peptides from rat tissue were separated on an Aurora column with captive spray insert (C18 1.6 uM particles, 25 cm, 75 μm inner diameter; IonOptics) at 60 °C. A solvent gradient of buffer A (0.1 % formic acid) and buffer B (99.9% acetonitrile/0.1% formic acid) over 140 min, at a flow rate of 600 nL/min, were used, respectively. The mass spectrometer was operated in DIA PASEF mode with 1.1 s cycle time and TIMS ramp time of 100.0 ms. MS scan range was set to 100–1700 m/z.

### Protein identification by mass spectrometry

Raw data-independent acquisition (DIA) files were analyzed with DIA-NN software (v.1.8.0) using UniProt FASTA database (UP000002494_10116.fasta (21,587 entries) and UP000002494_10116_additional.fasta (9981 entries), August 2020) and deep learning-based spectra to generate a library. In DIA-NN, the options ‘FASTA digest for library-free search/library generation’ and ‘Deep learning-based spectra, RTs and IMs prediction’ were enabled and all other settings were left default: Digestion protease = Trypsin/P; missed cleavage = 1; max number of variable modifications = 0; N-term M excision = True; C carbamidomethylation = True; peptide length range = 7-30; precursor charge range = 1-4; precursor m/z range = 300-1800; fragment ion m/z range = 200-1800; Precursor FDR (%) = 1%; Use isotopologues = enabled; MBR = enabled; Neural network classifier = single-pass mode; Protein inference = Genes; Quantification strategy = Any LC (high accuracy); Cross-run normalisation = RT-dependent; Library generation = Smart profiling; Speed and RAM usage = Optimal results.

MS raw files from a previous publication^7^ were download from the ProteomeXchange Consortium via PRIDE with the identifier PXD026051. The files contained DIA-MS information about mesenteric and renal arteries from 12-week-old SHR and WKY. The DIA-NN software was used for protein identification with the same settings as listed above.

### Bioinformatic analysis

Number of unique proteins at 1% FDR were obtained from the report file generated by DIA-NN. Protein quantities were obtained from the unique gene list in DIA-NN (*report*.*unique_genes_matrix*.*tsv*) and implemented in Perseus (v1.6.14.0)^16^. Data was log2 transformed and filtered for by 3 valid values in at least one group. Statistical comparisons between 12-week old rat groups was made using the limma package for R^17,18^ to enable batch correction and calculation of statistical significance levels and fold change difference per protein. Furthermore, two-sided Student’s *t*-test in Perseus was used to compare PFC groups, pre-hypertensive groups as well as previously published mesenteric and renal artery data^7^, with permutation-based FDR (<0.05) and 250 randomizations. Missing values were only imputed for PCA (width =0.4, down shift = 1.8). ECM and angiogenesis enrichments were achieved by comparing with a curated gene list^19–22^ and selecting overlapping proteins for further analysis. The pro- or anti-angiogenic property of angiogenesis-associated proteins was manually determined by literature mining. Hierarchical clustering was created based on z-scored LFQ values and generated by average linkage, preprocessing with k-means, and Euclidean distance. The z-score normalization was calculated by subtracting mean intensity from each protein value across all samples followed by division by the standard deviation^7^. ClueGO (v2.5.8)^23^ in Cytoscape (v3.9.1)^24^ was used to generate protein-protein interaction networks and enrichment analysis of significantly regulated proteins. The enrichment analysis was performed as previously described^7^; Rattus norvegicus was selected as organism, and the Gene Ontology (GO) biological processes (GO-BiologicalProcess, CellularComponent, ImmuneSystemProcess, MolecularFunction-EBI-UniProt-GOA-ACAP-ARAP, downloaded 15.01.2021) and Kyoto Encyclopedia of Genes and Genomes (KEGG, downloaded 15.01.2021) were used as ontology reference sets. The following settings were used; GO Tree interval = 3-8; GO Term/Pathway selection = 3 min #Genes, 4% genes; Kappa-score = 0.4; Enrichment/Depletion (Two-sided hypergeometric test with Benferroni step down, *p* ≤ 0.05; GO term fusion was enabled.

## Results

### Cerebral arteries in early-onset hypertensive rats display increased wall remodeling

The basilar artery was dissected in a subset of 12 week old rats (n=4), and stained with Sirius red to investigate the arterial morphology at a macroscopic level (Fig. 1a, 1b). This confirmed a significantly increased media diameter (Fig. 1c) and a reduced lumen diameter (Fig. 1d), accompanied with significantly increased media-to-lumen ratio in the 12-week-old SHRs compared to WKY controls (Fig. 1e).

**Figure 1:**
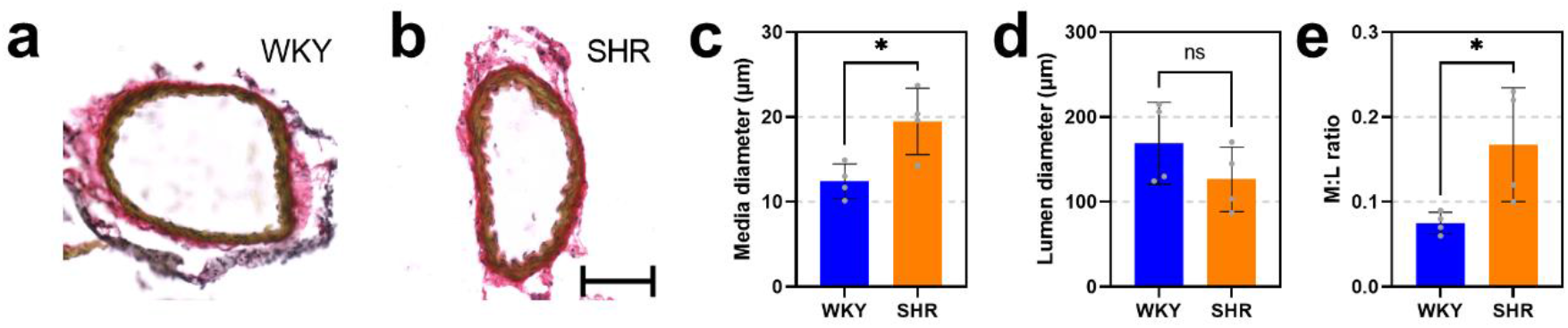
Vascular remodeling in cerebral arteries. a-b) Exemplary Sirius red stained basilar arteries from a subgroup of 12-week-old WKY (a) or SHR (b) rats (20× lens; scale bar = 100 μm). c-d) Media and lumen diameters in arterial cross-sections (n=4). e) The media-to-lumen (M:L) ratio was significantly higher in cerebral arteries from the 12-week-old SHR compared to WKY. Two-way Student’s *t*-test. *, *p* < 0.05. WKY = blue, SHR = orange.

### Cerebral arteries in hypertensive rats show distinct proteomic profile compared to normotensive control

After confirming a structural difference in cerebral vasculature of the SHR compared to the WKY, we investigated the protein composition by label-free DIA-MS quantification. We analyzed SHR and WKY cerebral arteries at 12 weeks of age (n=12) to capture the protein changes occurring in the early-onset hypertensive state (Fig. 2a). We identified a total of 4965 unique proteins that were observed across the cerebral artery samples, supporting the sensitivity of the DIA-MS approach with high reproducibility between samples (Fig. 2b and 2d). Using unbiased principal component analysis (PCA), we observed two clusters corresponding to phenotype (SHR and WKY) along component 2, accounting for 8.3% variation (Fig. 2c).

**Figure 2:**
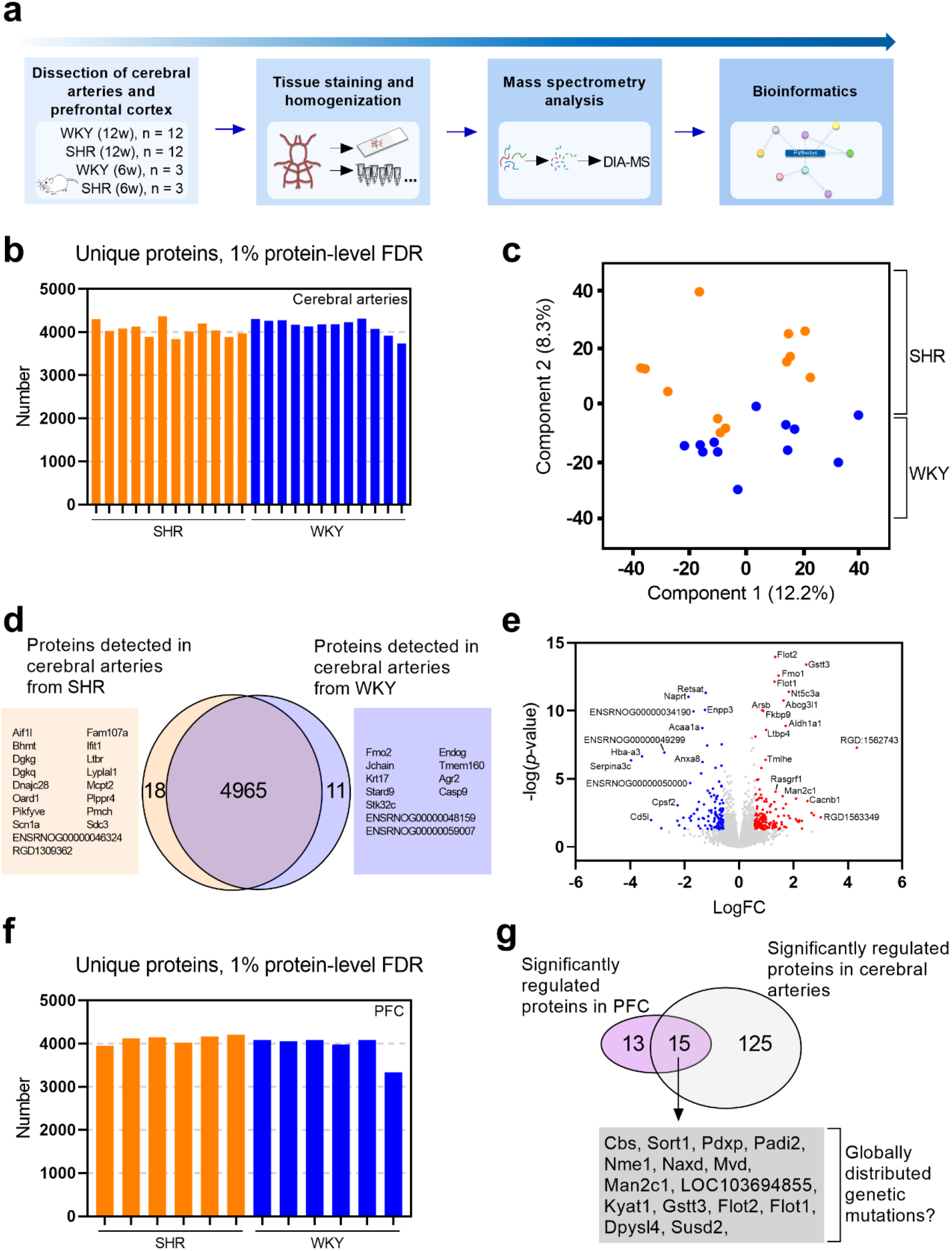
Identification of significantly regulated proteins in cerebral arteries from hypertensive rats. a) Study overview of the proteomic analysis. Cerebral arteries were dissected from two cohorts of Wistar Kyoto (WKY) and spontaneously hypertensive rats (SHR) at 6 week (n=3) and 12 weeks of age (n=12). A segment of the prefrontal cortex (PFC) was extracted from the 12-week-old rats (n=6). Extracted tissue was snap frozen and analyzed by histological staining or mass spectrometry (MS) analysis. Digested samples for MS analysis were analyzed by liquid chromatography–tandem MS (LC-MS/MS) using data-independent acquisition (DIA). Protein identification and quantification was obtained using DIA-NN software. b) Stacked bar representation of unique proteins identified by DIA-MS across cerebral artery samples from 12-week-old SHR (orange) and WKY (blue). FDR = false-discovery rate. c) Principal component analysis (PCA) plot of log2 transformed intensities associated with the 12-week-old rat samples. Components 1 and 2 are presented. d) Venn diagram showing total number of exclusive and shared proteins identified in cerebral arteries from 12-week-old SHR and WKY (orange and blue circle, respectively). e) Volcano plots comparing protein abundance in cerebral arteries from 12-week-old SHR and WKY controls. Log2 fold change difference and non adjusted *p*-value are presented in volcano plot. Red = upregulated and blue = downregulated in SHR compared to WKY. f) Stacked bar representation of unique proteins identified by DIA-MS across medial PFC samples from 12-week-old SHR (orange) and WKY (blue). g) Venn diagram representing number of significantly regulated proteins in statistical analysis of cerebral arteries and PFC from 12-week-old SHR and WKY rats. The number of shared proteins (n=15) were subsequently removed from the cerebral artery list because of a possible global genetic involvement.

Using a linear model (modified Student’s *t*-test)^17^, we identified 168 up- and 121 downregulated proteins (volcano plot visualization in Fig. 2e). After correcting *p*-values for multiple comparisons, we reduced the list to 140 proteins (73 up- and 67 downregulated), where expression levels were significantly different in the cerebral arteries from the SHR and WKY controls.

The SHR was derived from the WKY and inbred to perpetuate the hypertensive phenotype. The elevated BP is therefore based on several mutations^6^, many of which are likely to be unknown. To limit the influence of this factor, we performed an additional DIA-MS analysis of brain tissue samples from the medial prefrontal cortex (PFC) region of SHRs and WKY controls, and observed a similar unique protein identification (∼4000 proteins per sample) (Fig. 2f). Using a linear regression model, we identified 28 significantly regulated proteins between SHR and WKY controls in the PFC region. Of these, 15 proteins were also present in the analysis of cerebral arteries (Fig. 2g). These overlapping proteins may be caused by genetic mutations with a global distribution affecting different tissues in the SHR model, which is not specific to the vasculature. Based on the scope of this study, we therefore removed the 15 overlapping proteins, which left us with 125 significantly regulated proteins in the cerebral arteries from SHRs compared to WKY controls (Table 1; Fig. 2g).

**Table 1:**
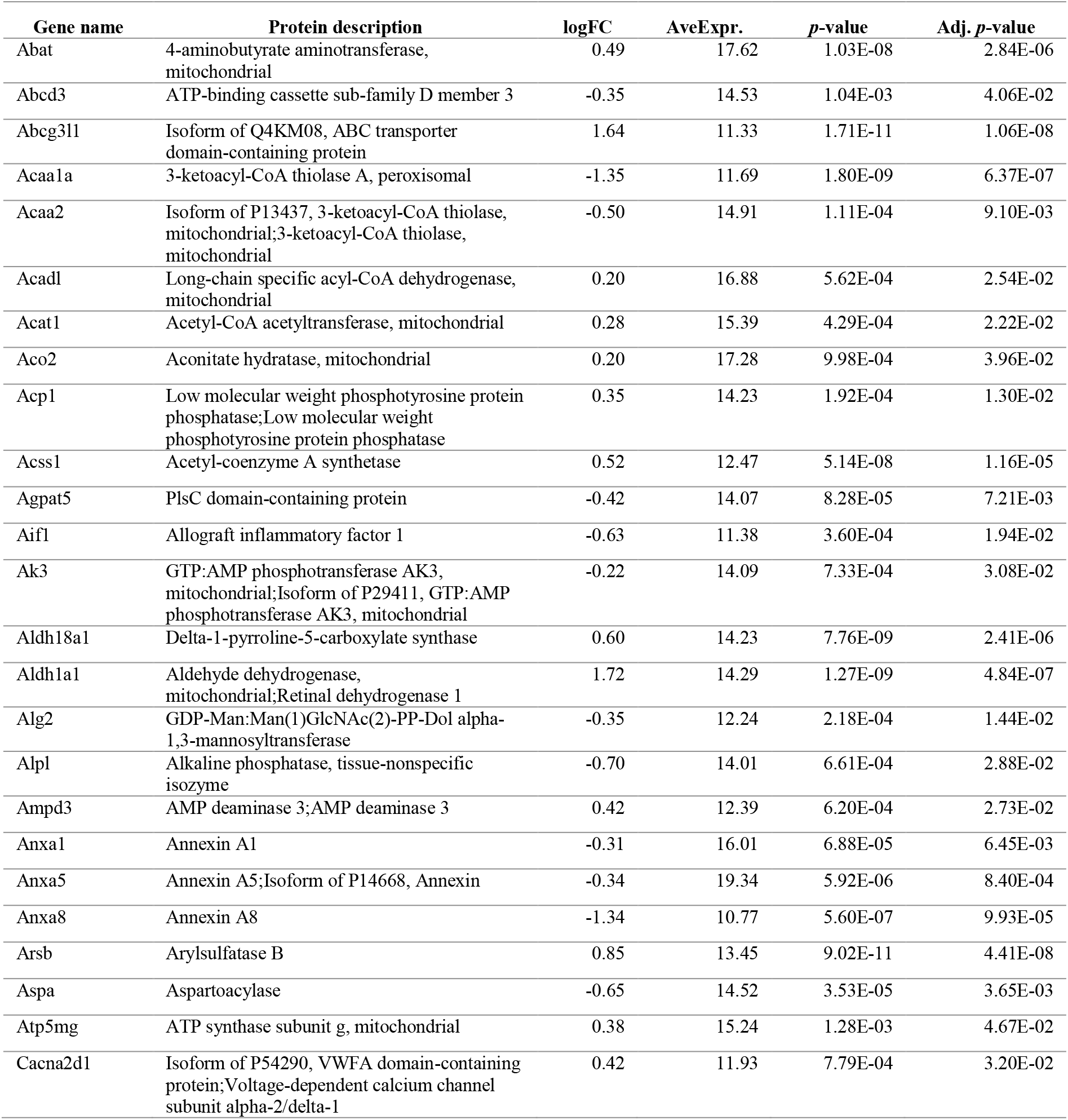
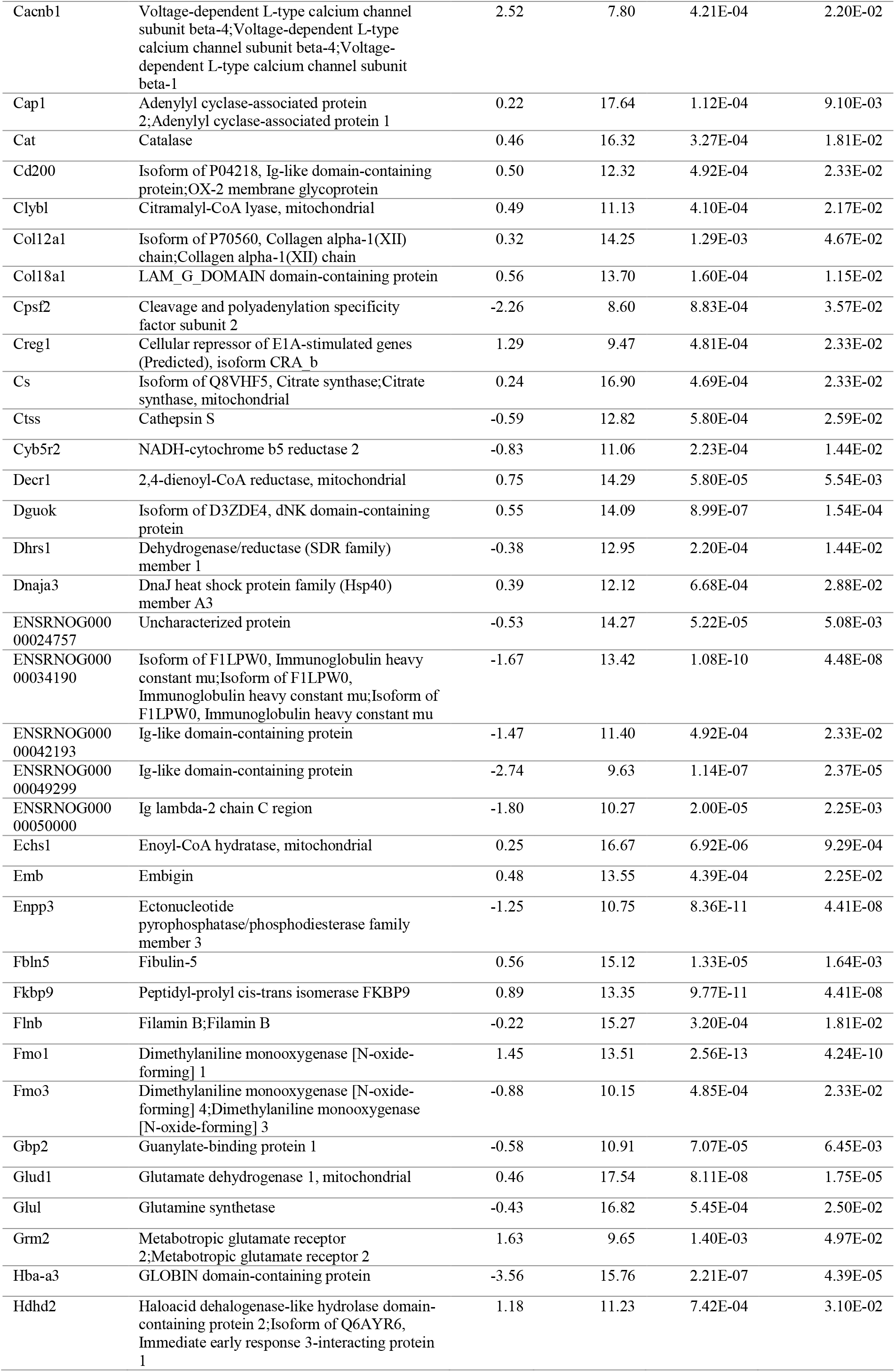
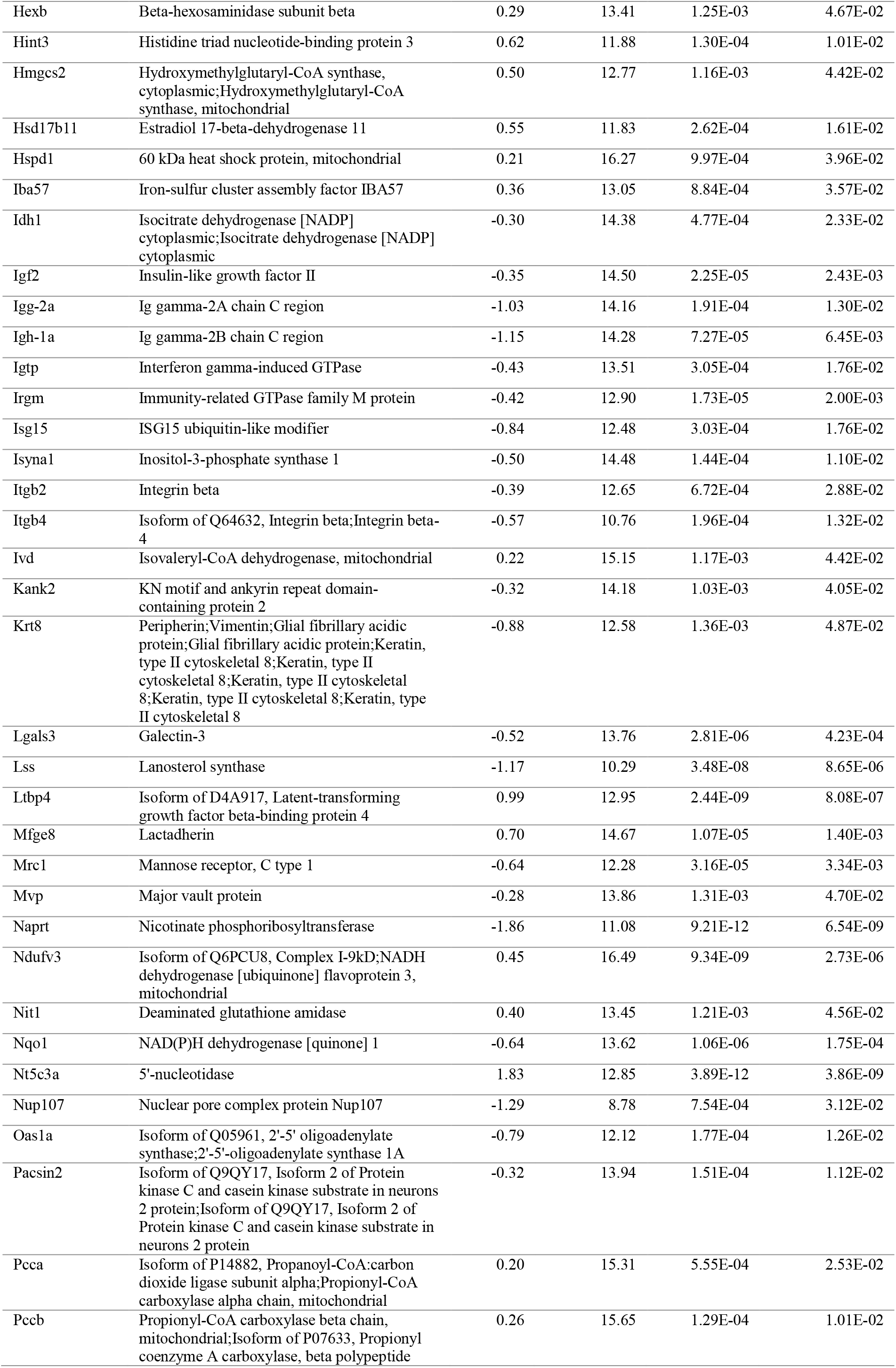
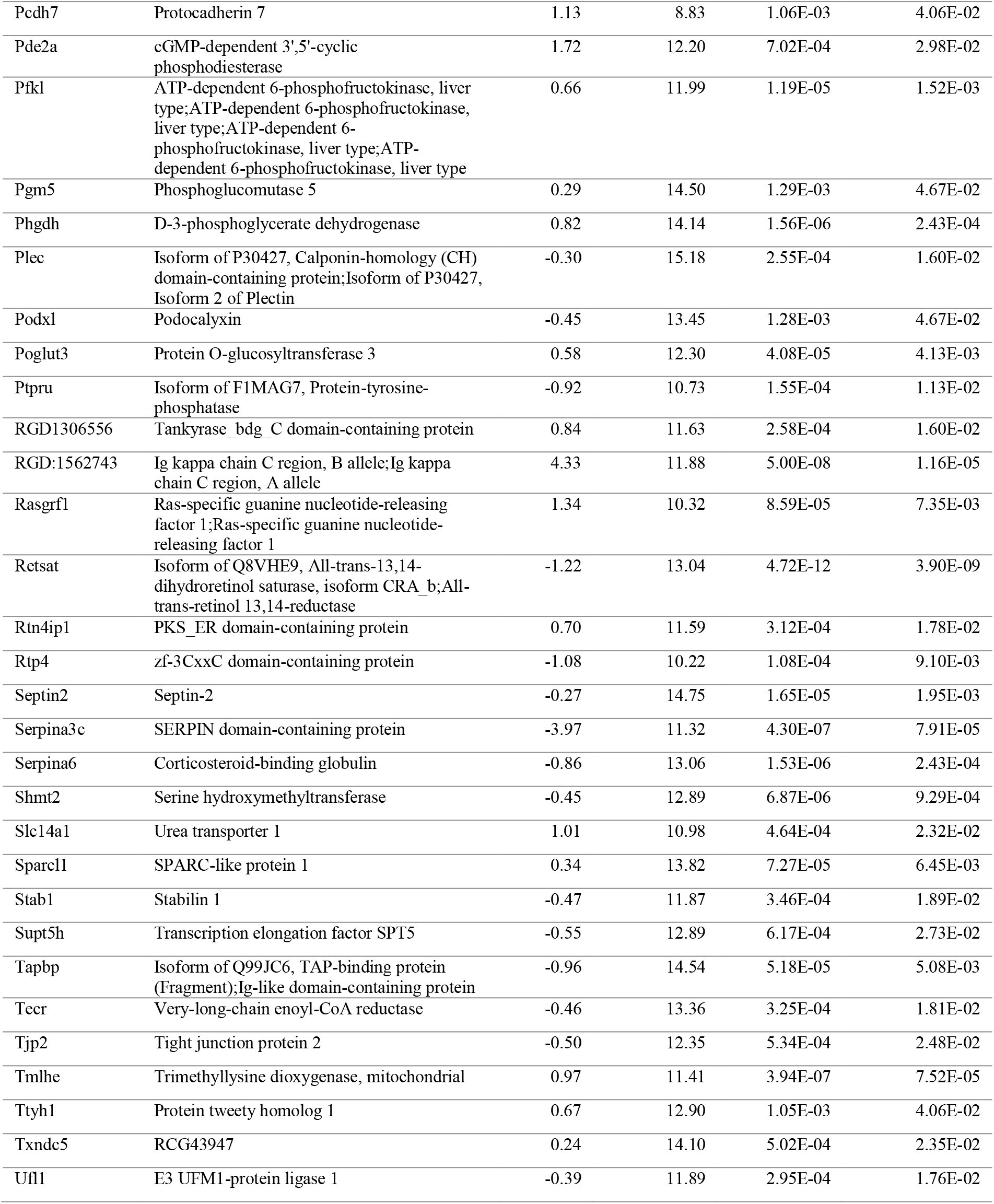
List of significantly regulated proteins detected in cerebral arteries from hypertensive rats compared to normotensive control. Description of the significantly regulated proteins identified when comparing the proteome of 12-week-old cerebral arteries from spontaneously hypertensive rats (SHR) and normotensive Wistar Kyoto rats (WKY). logFC =log2 fold change difference. AveExpr. = average expression. SHR vs WKY comparison.

### Pathway analysis identifies mechanistic differences in hypertensive rats compared to normotensive control

We next investigated whether particular pathways were regulated in the SHR compared to the WKY control. Using gene overrepresentation analysis, we identified 14 clusters of related Gene Ontology (GO) and Kyoto Encyclopedia of Genes and Genomes (KEGG) terms (Fig. 3a; Table 2). The clusters with lowest group term significance score were “Glyoxylate and dicarboxylate metabolism”, “Valine, leucine and isoleucine degradation”, and “fatty acid catabolic process” (Fig. 3b).

**Table 2:** Pathway analysis of proteins associated with hypertension. ClueGO-enrichment analysis of significantly regulated proteins identified in cerebral arteries from the comparison of 12-week-old spontaneously hypertensive rats (SHR) and normotensive Wistar Kyoto rats (WKY). The annotations represent related Gene Ontology (GO) or Kyoto Encyclopedia of Genes and Genomes (KEGG) terms enriched as nodes. The term represent the highest significance score.

| Term | Ontology Source | Term *p*-value corrected with Bonferroni step down | Group *p*-value corrected with Bonferroni step down | % Associated Genes | Associated Genes Found |
| --- | --- | --- | --- | --- | --- |
| negative regulation of cytokine-mediated signaling pathway | GO Biological Process | 6.18E-03 | 6.18E-03 | 4.7 | Dnaja3, Isg15, Oas1a |
| phagocytosis, engulfment | GO Biological Process | 1.10E-02 | 2.75E-03 | 5.4 | Aif1, Igh-6, Itgb2, Mfge8 |
| response to hydrostatic pressure | GO Biological Process | 8.57E-04 | 1.84E-04 | 27.3 | Col18a1, Krt8, Plec |
| vitamin metabolic process | GO Biological Process | 1.15E-02 | 4.61E-03 | 4.2 | Alpl, Clybl, Nqo1, Shmt2 |
| cellular response to interferon-beta | GO Biological Process | 1.43E-03 | 3.10E-04 | 10.8 | Gbp2, Igtp, Irgm, Oas1a |
| hydrolase activity, acting on carbon-nitrogen (but not peptide) bonds | GO Molecular Function | 3.62E-03 | 7.61E-04 | 4.2 | Ampd3, Aspa, Cat, Cd200, Hint3, Nit1 |
| cellular amino acid catabolic process | GO Biological Process | 9.68E-05 | 2.42E-07 | 6.6 | Abat, Acat1, Glud1, Ivd, Pcca, Pccb, Shmt2 |
| steroid biosynthetic process | GO Biological Process | 1.38E-03 | 8.03E-05 | 4.2 | Abcd3, Cyb5r2, Hmgcs2, Hsd17b11, Igf2, Lss, Tecr |
| acetyl-CoA C-acetyltransferase activity | GO Molecular Function | 2.11E-04 | 1.81E-04 | 42.9 | Acaa1a, Acaa2, Acat1 |
| Glyoxylate and dicarboxylate metabolism | KEGG | 4.53E-12 | 2.88E-13 | 29.0 | Acat1, Aco2, Acss1, Cat, Cs, Glul, Pcca, Pccb, Shmt2 |
| nucleoside phosphate catabolic process | GO Biological Process | 1.82E-03 | 4.21E-05 | 6.5 | Acat1, Ampd3, Enpp3, Nt5c3a, Pde2a |
| cellular amino acid biosynthetic process | GO Biological Process | 1.19E-05 | 4.52E-11 | 9.1 | Abat, Aldh18a1, Aldh1a1, Glud1, Glul, Phgdh, Shmt2 |
| Valine, leucine and isoleucine degradation | KEGG | 1.47E-09 | 1.21E-11 | 16.1 | Abat, Acaa1a, Acaa2, Acat1, Echs1, Hmgcs2, Ivd, Pcca, Pccb |
| fatty acid catabolic process | GO Biological Process | 2.19E-09 | 1.21E-11 | 9.8 | Abcd3, Acaa1a, Acaa2, Acadl, Acat1, Aldh1a1, Decr1, Echs1, Ivd, Pcca, Pccb |

**Figure 3:**
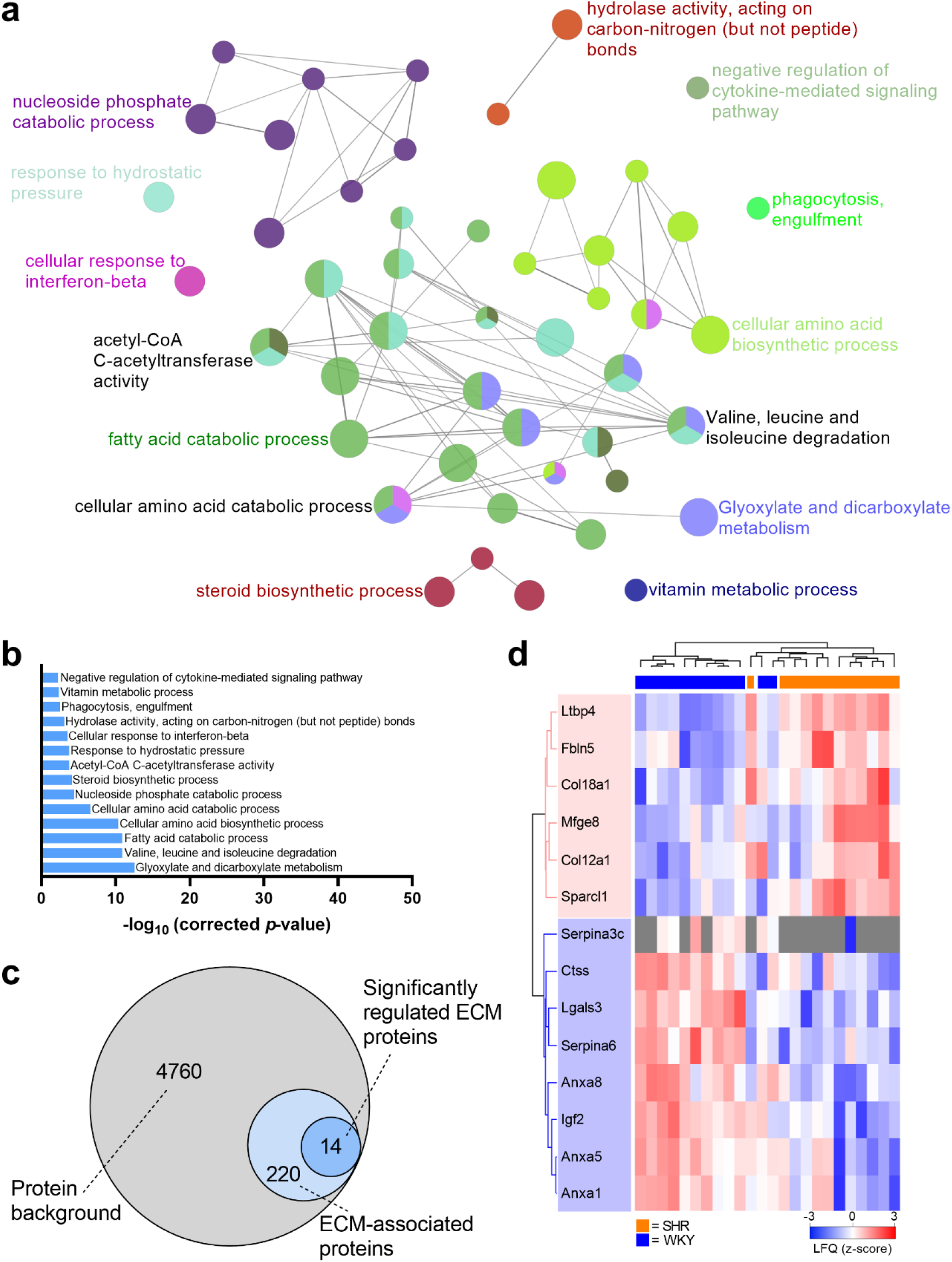
Pathway and clustering analysis of proteins identified in cerebral arteries from hypertensive rats. a) Gene overrepresentation analysis of significantly regulated proteins (n = 125) identified from the spontaneously hypertensive rats (SHR) vs normotensive Wistar Kyoto rats (WKY) comparison using ClueGO. The size of each node is based on a significance score and each cluster of related terms is labeled with the highest significance score of annotated ontology reference sets after *p*-value correction. b) Significance score identified in enrichment analysis of regulated proteins in SHR compared to WKY. c) Venn diagram showing the number of extra cellular matrix (ECM)-associated proteins in dataset. d) Unsupervised hierarchical clustering of 14 significantly regulated ECM-associated proteins identified in (c). Z-scored label-free quantification (LFQ) values from cerebral artery samples are depicted. SHR = orange, WKY = blue.

We have previously identified significant changes in the ECM-associated proteins in mesenteric arteries from 12-week-old SHRs^7^. Based on this previous finding, we enriched our cerebral artery data for ECM proteins using the same ECM database^7,19,20^. We identified 234 ECM-associated protein in our total dataset, 14 of which were regulated significantly between SHR and WKY (Fig. 3c). However, using unsupervised hierarchical clustering analysis, we did not observe a clear grouping of SHR and WKY based on protein expression intensities (z-score normalized; Fig. 3d), suggesting that the significantly regulated ECM proteins alone were not able to differentiate the two groups.

### Early-onset hypertensive rats display an imbalanced angiogenic profile in cerebral arteries but not in other systemic arteries

Hypertension and many cerebrovascular diseases are associated with compromised vascular integrity and promotion of vascular rarefaction. To investigate the latter, we compared our total protein background in the cerebral arteries to an angiogenesis database (GO term enrichment) because dysregulated angiogenesis can lead to vascular rarefaction (loss of capillaries and arterioles)^1^. This enrichment identified 184 proteins related to angiogenesis. Using a PCA plot based on the angiogenesis-associated proteins only, we observed two clusters corresponding to the phenotype (SHR and WKY) along component 1, accounting for 36.6% variation (Fig. 4a). This clustering was supported by an unsupervised hierarchical clustering analysis of significantly different angiogenesis-associated proteins (non-adjusted *p*-value) that similarly showed clear grouping of SHR and WKY based on protein expression intensities (z-score normalized; Fig. 4b).

**Figure 4:**
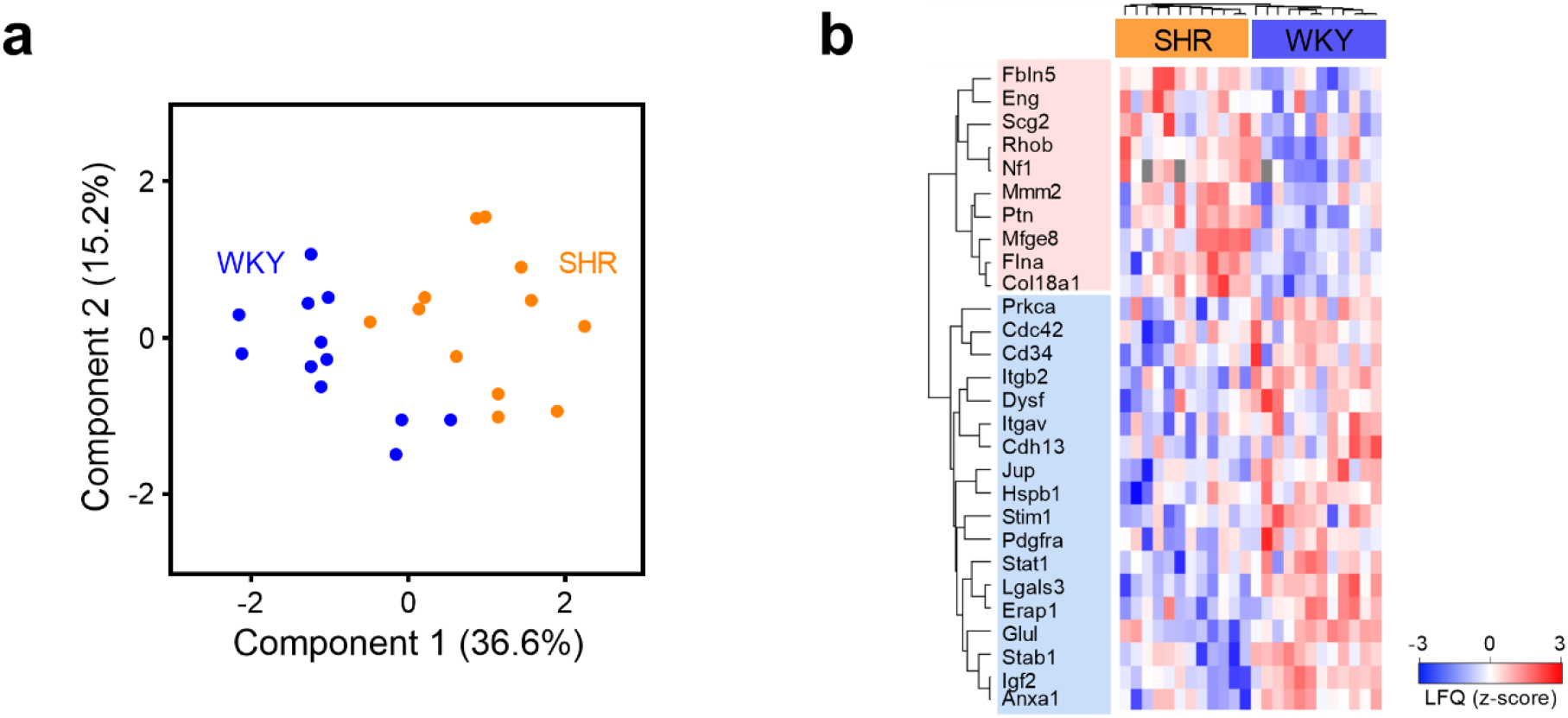
Clustering analysis of angiogenesis-associated proteins can differentiate between cerebral arteries from hypertensive and normotensive rats. a) Principal component analysis (PCA) plot of log2 transformed intensities associated with angiogenesis-assocaited proteins from 12-week-old spontaneously hypertensive rats (SHR, orange) or Wistar Kyoto rats (WKY, blue). Components 1 and 2 are presented. b) Unsupervised hierarchical clustering of significantly regulated angiogenesis-associated proteins (Two sided Student’s *t*-test, 75% valid values in at least one group). Z-scored label-free quantification (LFQ) values are depicted.

Next, we used the fold change difference identified from the SHR vs WKY comparison (with a moderated *t*-test), to create a plot encompassing angiogenic changes (pro-angiogenic = red, anti-angiogenic = blue, unknown or both = green). Importantly, a clear pattern of upregulated anti-angiogenic proteins (=7/8, 87.5%), and downregulated pro-angiogenic proteins (17/24, 70.83%) was observed (Fig. 5a, 5b; Tables S1).

**Figure 5:**
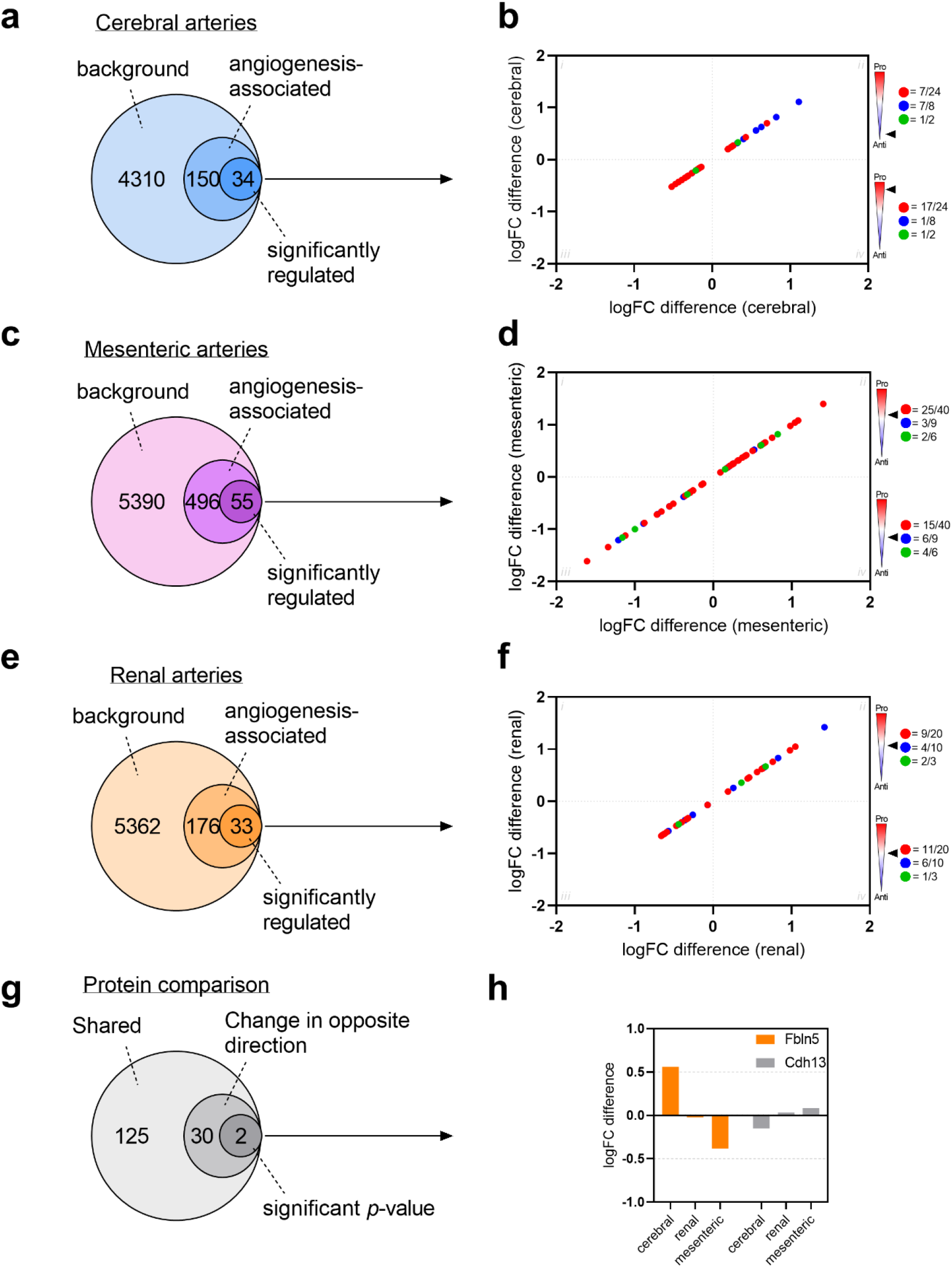
Angiogenic profile in three vascular beds from early-onset hypertensive rats. a) Venn diagram representing the number of angiogenesis-associated proteins in the cerebral artery dataset. The annotations were based on the Gene Ontology (GO) reference set (GO:0001525). b) Scatter plot of log2 fold changes (logFC) for all significantly regulated angiogenesis-associated proteins when comparing proteomic profile of cerebral arteries from 12-week-old spontaneously hypertensive rats (SHR) and normotensive Wistar Kyoto rats (WKY) (moderated *t*-test; non-adjusted *p*-value was used). Each protein is colored as pro-angiogenic (=red), anti-angiogenic (=blue), and unknown or both (=green). Black arrow indicates angiogenic profile in respective squares (ii-iii). c,e) Venn diagrams representing the number of angiogenesis-associated proteins in the mesenteric and renal artery datasets from a previous publication^7^. d,f) Scatter plot similar to (b) representing fold change differences of angiogenic proteins in mesenteric and renal arteries from SHR and WKY comparisons, respectively. g) Venn diagram showing number of angiogenesis-associated proteins in cerebral arteries that are shared with mesenteric and renal arteries, but change in opposite direction. h) Column graph depicting logFC difference of Fbln5 (=orange) and Cdh13 (=grey) in cerebral, renal and mesenteric artery comparisons between 12-week-old SHR and WKY.

To determine whether this angiogenic pattern was maintained in different arterial beds, we applied the same analysis to our recently published database on mesenteric and renal arteries in SHRs at the early-onset stage. To enable this comparison we re-analyzed the data with the same analysis software as presented in this study (DIA-NN). This led to a total identification of 5941 and 5571 unique proteins in the mesenteric and renal arteries, respectively (n=14 samples in each arterial bed type (SHR=7 and WKY=7)). After comparing both datasets to the angiogenesis database (GO term enrichment) and applying a *p*-value cutoff <0.05 (Student’s *t*-test), we identified 55 and 33 significantly regulated angiogenesis-associated proteins in the mesenteric and renal artery datasets, respectively (non-adj. *p*-values; Fig. 5c, 5e). When creating similar plots encompassing protein expression differences, we observed that both pro- and anti-angiogenic proteins were distributed equally between up- and downregulation (Fig. 5d, Fig. 5f; Tables S2 and S3). This suggested that the angiogenic imbalance observed in the cerebral arteries was not systemic but rather local to the cerebrovasculature.

Following this, we investigated the protein changes that were potentially driving the observed angiogenic difference. Thus, we identified angiogenic proteins that were shared between the three vascular beds (before *p*-value filtering), and focused on proteins expressed in the cerebral arteries that were changing in opposite direction to mesenteric and renal arteries (Fig. 5g). This left us with 32 proteins, of which two (Fbln5 and Cdh13) were significantly regulated in the cerebral artery comparison from SHRs and WKY controls (Fig. 5h).

### Imbalanced angiogenesis is not present in pre-hypertensive rats

We investigated the protein composition in cerebral arteries from pre-hypertensive rats (6-week-old) and age-matched WKY controls by label-free DIA-MS quantification (n=3 in each group). Our approach led to a reproducible identification across the 6-week-old samples in both groups (Fig. 6a). When comparing to the 12-week-old dataset, we identified 4673 shared proteins that were observed in both datasets, supporting similar proteomic backgrounds despite the age difference (Fig. 6a and 6b).

**Figure 6:**
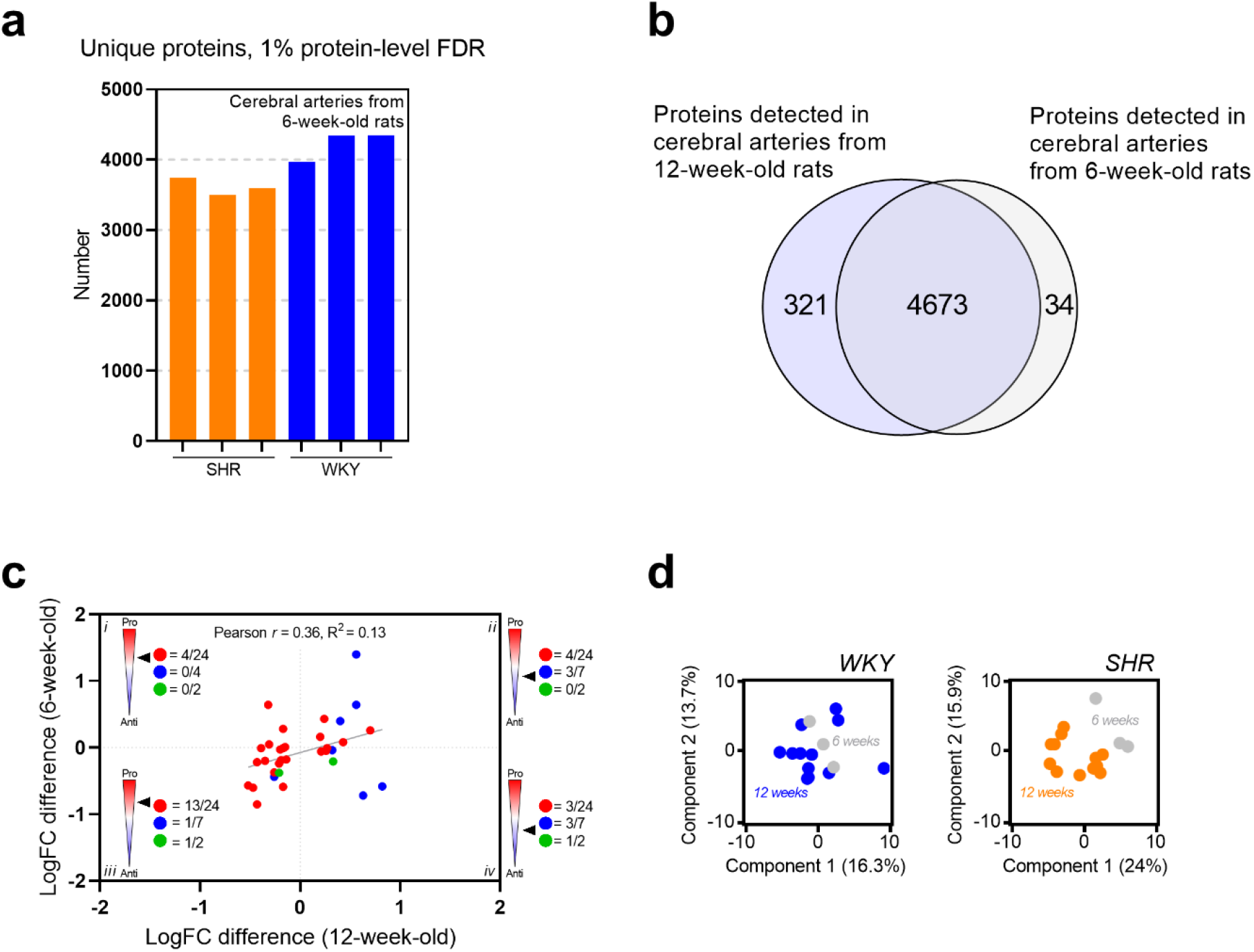
Blood pressure-induced angiogenic imbalance in the hypertensive rat. a) Stacked bar representation of unique proteins identified by data-independent acquisition mass spectrometry (DIA-MS) across cerebral artery samples from 6-week-old spontaneously hypertensive rats (SHR, orange) and Wistar Kyoto (WKY, blue). b) Venn diagram depicting protein overlap between 6- and 12-week-old cerebral artery datasets. c) Scatter plot of log2 fold changes (logFC) for all significantly regulated angiogenesis-associated proteins when comparing proteomic profile of cerebral arteries from 6- and 12-week-old spontaneously hypertensive rats (SHR) and normotensive Wistar Kyoto rats (WKY) (x-axis = 12-week-old, y-axis = 6-week-old). Each protein is colored as pro-angiogenic (=red), anti-angiogenic (=blue), and unknown or both (=green). Black arrow indicates angiogenic profile in respective squares (i-iv). Pearson *r* coefficient (= 0.36) is indicated by grey line. d) Principal component analysis (PCA) plot of log2 transformed intensities from angiogenesis-associated proteins associated with WKY samples (left) and SHR samples (right). Components 1 and 2 are presented.

To determine whether the angiogenic changes identified in 12-week-old cerebral arteries from SHRs were also present at the pre-hypertensive state, we performed a similar angiogenic enrichment of shared proteins. We then created correlation plots using the expression difference calculated between SHR vs WKY at both time points (Fig. 6c). Both pro- and anti-angiogenic proteins were upregulated at both the 6- and 12-week comparisons (Fig. 6c; Tables S4). Furthermore, the Pearson *r* and R^2^ coefficients (0.36 and 0.13, respectively; Fig. 6c) did not support a correlation between the two time-points.

To account for age-associated changes within the WKY or SHR groups independently, we applied PCA plots on shared angiogenesis-associated proteins only (Fig. 6d). No clustering was observed in the 6- and 12-week-old WKY samples, suggesting that aging alone was not capable of changing the angiogenic profile. On the contrary, a clear clustering association was observed when looking at the 6- and 12-week-old SHR samples on component 1, suggesting that the angiogenic imbalance detected in the cerebral arteries was driven by elevated blood pressure in the SHR model, and thus blood pressure dependent (Fig. 6d).

Interestingly, expression of Fbln5 was upregulated significantly in the 6-week old SHRs compared to WKYs (logFC difference = 0.64; *p*-value = 0.033, Adj. *p*-value = 0.411), whereas the expression of Cdh13 was not different between the 6-week old SHR and WKY control (logFC difference = 0.01; *p*-value = 0.959, Adj. *p*-value = 1).

## Discussion

Remodeling of the arterial wall in hypertension is a critical determinant of disease progression and contributes to the risk of developing cardiovascular disease. In order to identify the pathways and mechanisms underlying hypertension-related vascular remodeling, we have employed an in-depth proteomic approach in this study and in our previous publication^7^. In this study, we identified several proteins with dysregulated expression in the cerebral vasculature of the SHR, early after the onset of hypertension (12-week old). Enrichment analysis revealed that proteins involved in angiogenesis were changed in the SHR cerebral arteries in a pattern suggestive of reduced angiogenesis, which was not observed in the resistance mesenteric or conduit renal arteries of age-matched SHRs. Furthermore, our experimental setup revealed two potential candidate proteins (Fbln5 and Cdh13), whose expressions were critically changed in cerebral arteries compared to systemic arteries at the early-onset of hypertension in the SHR.

In order to prevent neurovascular complications associated with hypertension, we need a better overview of the changes occurring in the cerebrovasculature at the initial stages of the disease. Our proteomic analysis identified 125 significantly regulated proteins that were changed the early-onset of hypertension in the SHR model. Our enrichment analysis of these proteins revealed that *Glyoxylate and dicarboxylate metabolism* was the most predominant pathway affected. Glyoxylate is the conjugant base of glyoxylic acid and involved in pH regulation. Five of the associated proteins are also directly (Cs, Aco2) or indirectly (Acss1, Pcca, Pccb) involved in the citric acid cycle, and thus oxidation of carbohydrates and fatty acids. The protein expression of all five proteins was significantly upregulated in the cerebral arteries from the SHR compared to the WKY control at the early-onset of hypertension. A continued activation of the citric acid cycle together with upregulation of reactive oxygen species (ROS) homeostasis was observed in resistant hypertensive patients^25^. Excessive elevation of ROS can damage of the vascular wall leading to vascular remodeling through a multitude of pathwyas^26^. These protein and pathway occurred in 12-week-old SHR rats, supporting a critical mechanistic impact on cerebral arteries at the early-onset of hypertension.

Previously, we showed that ECM-related proteins were highly dysregulated in the resistance mesenteric arteries of the 12-week old SHR, but such changes were less prevalent in the SHR cerebral arteries, in line with a less pronounced change in the media-to-lumen ratio changes compared to the mesenteric arteries^7^. Our previous study found ECM changes correlated with significant hypertrophic remodeling with increased media-to-lumen-ratio of the arterial wall visualized by Sirius red staining. In our current study, we also detected a significantly increased media-to-lumen ratio in the cerebral arteries, which is consistent with previous observations in cerebral arterioles from stroke-prone SHR (SHRSP)^27^ and SHR^28^. However, the degree of remodeling was not as severe as the remodeling observed in the mesenteric arteries at the early-onset of hypertension (12-weeks old)^7^. The depressed impact of remodeling of cerebral arteries compared to systemic arteries in the SHR was also observed previously^13^, suggesting that the cerebral arteries are not remodeled to the same extent in early-onset hypertension. In line with this, we could not separate clearly the 12-week old SHR and WKY control when using unsupervised hierarchical clustering analysis based on the ECM-associated proteins. Furthermore, our analysis only detected 14 significantly regulated ECM-associated proteins compared to 38 proteins in mesenteric artery tissue in our previous study^7^.

Although structural arterial remodeling (inward eutrophic and hypertrophic) is an important factor known to contribute to increased total peripheral resistance in hypertension, vascular rarefication is another factor that can contribute to the increased resistance. Rarefaction can be due to fewer vessels connected in parallel, or due to a total closure of some of the vessels^29^. Vascular rarefaction is associated with the development of several clinical complications, such as cerebral microhaemorrhages and lacunar infarcts^30^. Reduced vascular density and angiogenesis has been demonstrated in the SHR^31^. The balance of pro- and anti-angiogenesis proteins controls the dynamic ability of arterioles to grow, recede or maintain their structure, thereby controlling vascular rarefaction^1^. In this study, we have identified key angiogenic proteins that are dysregulated in the cerebral arteries of the SHR. By assigning the proteins as either pro- or anti-angiogenic, we show a clear pattern indicating that angiogenesis is likely to be impaired in the cerebral arteries of the SHR, which would promote vascular rarefication. Importantly, this pattern was not observed in the resistance mesenteric or conduit renal arteries of the SHR, nor was it observed in 6-week old SHR cerebral arteries. Thus, early in the onset of hypertension, when systemic vessels undergo more ECM-dependent inward remodeling, the cerebral arteries display a shift towards reduced angiogenesis, a precursor for rarefaction.

Our experimental design allowed us to reveal two candidate proteins (Fbln5 and Cdh13) that were differentially expressed in the cerebral arteries compared to two systemic vascular beds at the early-onset of hypertension in the SHR. Fbln5 (Fibulin 5 or DANCE) is involved in the arrangement of elastic fibers in the vessel wall and has been reported to suppress the matrix metallopeptidase MMP9^32^. Mice that lack Fbln5 display fragmented and disorganized elastic fibers^33^. In contrast, overexpression of Fbln5 was found to increase elastin deposition in retinal pigment epithelial cells^34^. Furthermore, overexpression of Fbln5 in mice subjected to ischemia/reperfusion injury after middle cerebral artery occlusion, was found to attenuate ROS production and reduce blood-brain barrier permeability and apoptosis^35^. The anti-angiogenic function of Fbln5 was demonstrated both *in vitro* and *in vivo*, possibly via increased thrompospondin-1 expression, antagonizing VEGF^165^ signaling or the α5β1 fibronectin receptor^36^. Interestingly, a phenome-wide association analysis of FBLN5 in the white British participants of the UK Biobank identified an intronic single nucleotide polymorphism (rs1049468267) within the FBLN5 gene that was associated with “cerebral atherosclerosis” (P = 9.9×10-7; effect size [beta value] = 88) and “occlusion and stenosis of precerebral arteries” (P=2.0×10-4; effect size [beta value] = 16). These data suggest a strong association of fibulin 5 with cerebral artery diseases, and our data suggest that this protein is dysregulated in cerebral arteries early in hypertension, which may increase the risk of certain cerebrovascular diseases related to hypertension.

The calcium-dependent cell–cell adhesion glycoprotein Cdh13 (also Cadherin 13 or T-cadherin) can work as receptor for low-density lipoproteins (LDL) and adiponectin, and has demonstrated potential navigating functions in migrating cells^37^. Its location was found to be restricted to caveolae in aortic smooth muscle cells where it interacted with Src, supporting that Cdh13 can facilitate intracellular signaling^37^. Genome-wide association studies (GWAS’s) have identified genetic variants in *CDH13* that are associated with coronary artery disease^38^ and blood pressure traits^39^. Furthermore, increased Cdh13 expression was found to promote tumor angiogenesis *in vivo*^40^. Taken together, the changes in expression of these candidate proteins have direct impacts on vascular remodeling and angiogenesis. However, the molecular pathway to these changes and whether they are connected should be investigated in future studies. Importantly, the regulation of both proteins are early mechanistic changes in the cerebrovasculature induced by elevated blood pressure. With further investigation, these proteins have the potential to be innovative therapeutic targets reducing the long-term risk of developing brain diseases associated with hypertension.

Hypertension is a major risk factor for vascular dementia and Alzheimer’s disease (AD). Accumulating evidence suggests that reduced cerebrovascular microcirculation plays a key role in the progression of (AD)^30,41^, a feature that is observed in hypertensive patients^42^. Furthermore, hypertension can cause vascular damage, neurovascular uncoupling, infarcts, and micro/macro bleeds, all favoring the pathogenesis of AD and therefore the risk of development^43^. AD is characterized by excessive accumulation and deposition of extracellular amyloid-beta (Aβ) and intracellular hyperphosphorylated tau protein^44,45^. Deposition of Aβ is present in the neurovascular unit and increased deposition leads to degeneration of small vessels and capillaries as well as impaired clearance^41^. The cerebral blood flow is decreased and cerebrovascular resistance is increased in both human AD and mice overexpressing the human amyloid precursor protein (APP) transgene bearing mutations that are associated with familial AD^46^. Furthermore, cerebral blood flow reduction was observed in preclinical AD and correlated with cognitive impairment^47^, supporting a critical role of the cerebrovasculature in the early phase of AD development. Importantly, SHRs were recently found to develop hallmarks of AD (Aβ plaques, intracellular hyperphosphorylated tau, and cognitive impairment^48–54^). Future studies should therefore investigate whether an anti-angiogenic protein profile in hypertensive patients can trigger cerebrovascular rarefaction, reduction in cerebral blood flow, and thereby lead to the increased risk of developing the pathological changes observed in AD.

This study has some limitations; first, because the cerebral arteries were rapidly dissected to prevent protein degradation, leftover material from the meninges cannot be excluded. Second, the study was limited to the circle of Willis and the primary cerebral arteries, meaning that the arteries have been mainly conduit arteries with lack of brain penetrating arteries. However, by using these arteries we diminished the impact of our first limitation by reducing the inclusion of proteins derived from brain cells e.g. neurons, glial cells and pericytes. Third, the study did not investigate the presence of rarefaction or blood flow changes because of the early phase of the disease. We consider that such changes would not be present at this time point, but rather develop over time. Future studies should therefore investigate these parameters in older SHRs that may yield a different picture of the vascular response to elevated blood pressure^14^.

In summary, this study has discovered critical proteomic differences in the cerebrovasculature of hypertensive rats. In-depth protein profiling has identified a hypertension-induced imbalance of angiogenic proteins in cerebral arteries that upregulate anti-angiogenic proteins and downregulate pro-angiogenic proteins. Collectively, these data reveal novel protein changes in the cerebrovasculature at the early-onset of hypertension that, over-time, could promote to cerebrovascular rarefaction, thereby increasing the risk of cerebrovascular diseases associated with hypertension. By mapping the hypertension-associated changes in the cerebral artery wall, this study reveals potential novel proteins and pathways that could be targeted to reduce the risk of developing cerebrovascular diseases associated with hypertension.

